# Unfolding of the chromatin fiber driven by overexpression of bridging factors

**DOI:** 10.1101/2020.07.29.224972

**Authors:** Isha Malhotra, Bernardo Oyarzún, Bortolo Matteo Mognetti

## Abstract

Nuclear molecules control the functional properties of the chromatin fiber by shaping its morphological properties. The biophysical mechanisms controlling how bridging molecules compactify the chromatin are a matter of debate. On the one side, bridging molecules could cross-link faraway sites and fold the fiber through the formation of loops. Interacting bridging molecules could also mediate long-range attractions by first tagging different locations of the fiber and then undergoing microphase separation. Using a coarse-grained model and Monte Carlo simulations, we study the conditions leading to compact configurations both for interacting and non-interacting bridging molecules. In the second case, we report on an unfolding transition at high densities of the bridging molecules. We clarify how this transition, which disappears for interacting bridging molecules, is universal and controlled by entropic terms. In general, chains are more compact in the case of interacting bridging molecules since, in this case, interactions are not valence-limited. However, this result is conditional on the ability of our simulation methodology to relax the system towards its ground state. In particular, we clarify how, unless using reaction dynamics that change the length of a loop in a single step, the system is prone to remain trapped in metastable, compact configurations featuring long loops.

## INTRODUCTION

Nuclear DNA in eukaryotic cells is scaffolded by histones and other proteins forming a fiber known as chromatin. A myriad of molecules, mainly proteins and ribonucleic acids (RNAs), regulate the morphological properties of the chromatin by selectively tagging and bridging specific loci of the genome [1–4]. Different conformations of the chromatin fiber result in different functional states. For instance, a change in the concentration of a bridging molecule can open or fold specific segments of the fiber, allowing or not polymerase enzymes to diffuse towards a promoter, and therefore silencing or activating particular genes.

Coarse-grained models have been employed to study the three-dimensional structure of the chromatin. A typical class of models uses bead-and-spring chains (see Fig. 1) in which monomers (which may represent a segment of chromatin containing multiple histones) interact through selective interactions [5–8] which could change as a result of chemical reactions triggered by epigenetic regulators [9]. On one side, such models are used to sample the physical properties of the system. Moreover, bead-and-spring models are also used to reconstruct the most likely conformation of the fiber compatible with experimental results in chromosome-capture techniques [10] measuring the euclidean proximity between different chromatin loci (as expressed by connectivity maps) [11, 12].

**Figure 1.**
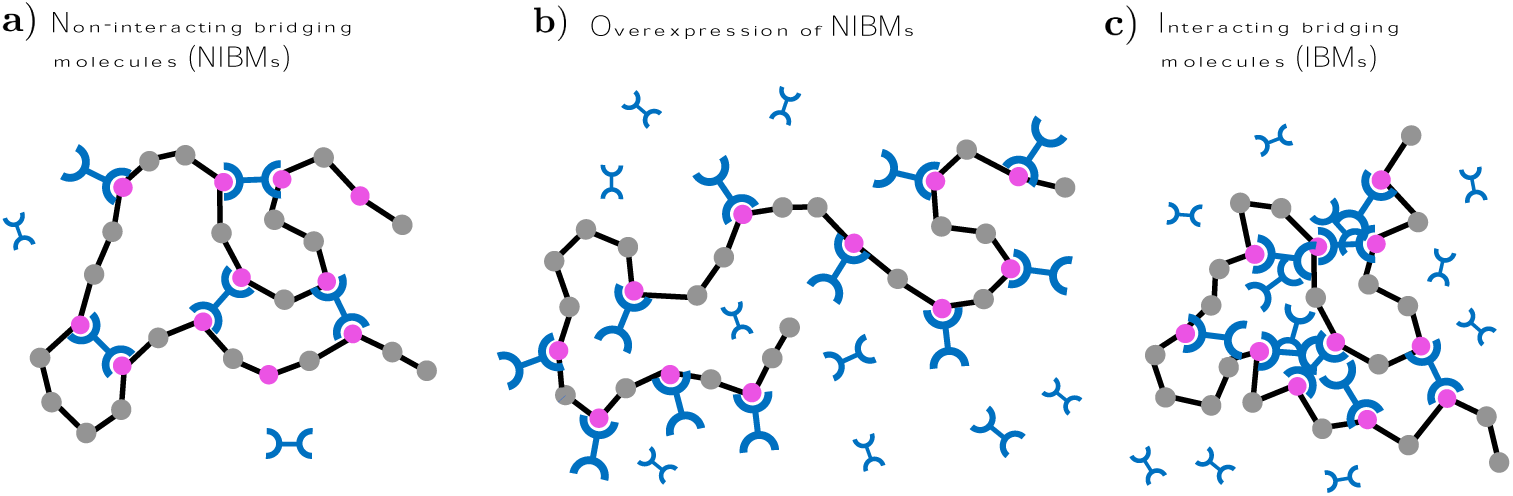
**a)** Non–interacting bridging molecules can bind tagged sites of the fiber (in purple) and drive compaction of the chromatin through the formation of loops. **b)** For high densities of the bridging molecules, each reactive monomer carries a B molecule, loops open, and the chromatin unfolds. **c)** Microphase separation of interacting B molecules can drive compaction of the fiber.

Simulation approaches are challenged by the multiscale nature of the system [13–15] as well by the necessity of sampling the large variability of the possible fiber configurations. In this paper, we address the latter issue and deploy an ensemble of Monte Carlo (MC) moves that allow changing the connectivity state of a chain in a single step by binding/unbinding two monomers while simultaneously regrowing a fraction of the chain. Using our simulation scheme, we study the morphological properties (compaction, connectivity map, and loop size distribution) of chromatin fibers cross-linked by bridging factors (see Fig. 1a). We study how the concentration of the bridging molecule *(ρ*_*B*_) and the affinity between bridging molecules and tagged (or reactive) monomers (quantified by the association constant, *K*_*a*_) affect the phase behavior of the system. Intriguingly, as foreseen in Ref. [16], for high values of *ρ*_*B*_ (and a given value of *K*_*a*_), the fiber unfolds given that all reactive monomers carry a linker, and bridges cannot form (see Fig. 1b). Simulation results are backed by scaling analysis and theory. The overexpression of a bridging factor can affect the phenotype (e.g. [17]). However, we are not aware of molecular studies of the chromatin fiber at high values of *ρ*_*B*_. The unfolding transition of Fig. 1b is somehow analogous to the resolubilization of virus crystals [18], proteins [19], or nucleic acids [20] at a concentration of multivalent ions (forming inter-molecular bridges) higher than some mMs. Similarly, aggregates of ligand-presenting colloids cross-linked by short DNA oligomers dispersed in solution melt when increasing the concentration of the bridging molecule beyond a threshold value (ranging between 10^*-*8^ M and 10^*-*4^ M) which depends on the number of ligands per particle [21]. The proposed method, along with other simulation strategies implementing reaction moves in systems of multivalent chains [22–30], are ideal for studying the unfolding transition given that the high free energy barriers between competing states are sidestepped by dedicated MC moves. On the other hand, MC methods are not suitable to study the dynamics of the system unless the reaction timescales are much bigger than the backbone relaxation times [27].

Loop formation is not the only way of compactifying chromatin’s segments. Bridging molecules may interact through residual (e.g., multivalent) interactions [31, 32]. Importantly, while remaining soluble in the solution, the bridging molecules may condensate on the chromatin fiber and form finite-sized drops enveloping reactive monomers (see Fig. 1c) [16]. The condensation of bridging molecules results in effective interactions between reactive monomers, which are not valence limited, finally driving the folding of the fiber [33]. In the presence of interacting bridging molecules, we find that the re-entrant transition disappears. In particular, the fiber remains in a compact form as more bridges are found onto the chain. This result only holds when comparing equilibrated states. When not changing the length of the loops with dedicated MC moves (which do not affect the equilibrium distribution of the system), chains folded by non-interacting bridges become much more compact. The results and methodology presented in this paper allow assessing the thermodynamics of competing mechanisms leading to domain formation in chromatin.

## METHODS

### The coarse-grained model

Fig. 2 presents the model. We consider fully flexible freely jointed chains made by *N* monomers with bond length equal to *σ. N*_*R*_ monomers out of *N* can reversibly bind bridging (B) molecules dispersed in solution. We evenly distribute the reactive (R) monomers along the chain and define the degree of functionalization as *f* = *N*_*R*_*/N*. *N*_1_ and *N*_2_ are, respectively, the number of R monomers carrying a bridge, not forming a loop, and the number of B molecules cross–linking two reactive monomers. In this study, we model bridging molecules implicitly without tracking their exact position in the system. Instead, after binding a bridge, an *R* monomer changes the way of interacting with the rest of the chain. Binding/unbinding events between B and R are directly controlled by the chemical potential of the B molecules *µ*_*B*_ and the association constant *K*_*a*_ (see below). For interacting bridges (see Fig. 1c), the attachment of B molecules to the fiber becomes cooperative.

**Figure 2.**
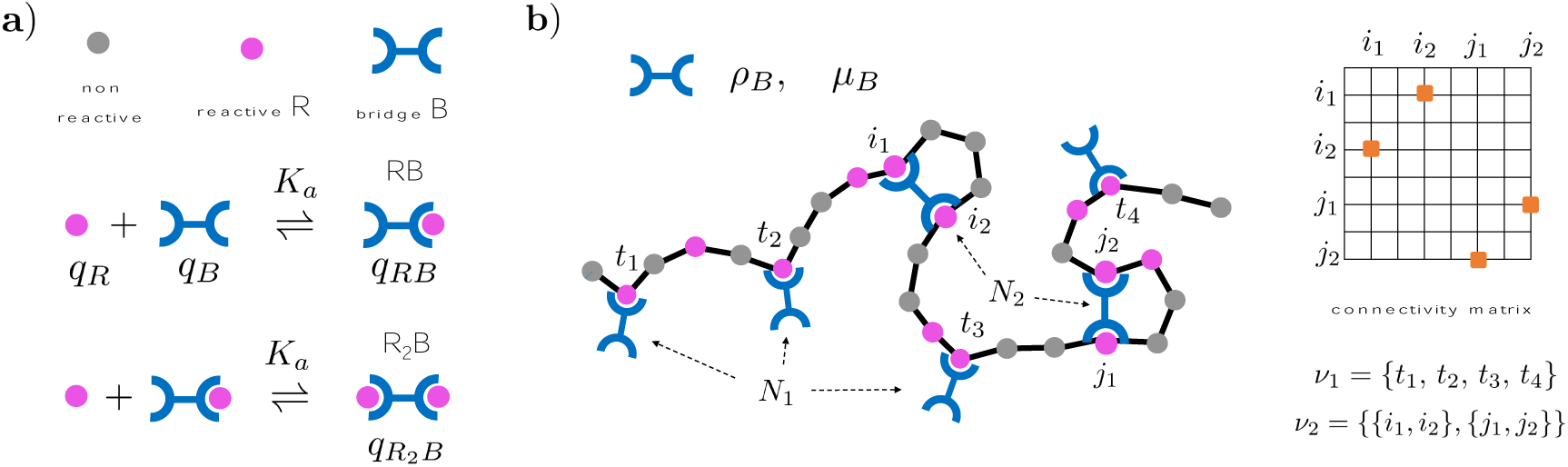
**a)** Formation of dimers (RB) and trimers (R_2_B) starting from reacting units (B molecules and R monomers) in diluted conditions. **b)** B molecules bind chains carrying R monomers and lead to the formation of loops. *µ*_*B*_ and *ρ*_*B*_ are, respectively, the chemical potential and the density of the B molecules in solution. *v*_1_ is the list of R monomers forming an RB complex while *v*_2_ the list of pairs of monomers cross-linked by B molecules. *v*_2_ can be represented using connectivity maps that, when averaged, provide the likelihood of finding two monomers cross-linked by a B molecule.

The microstates of the system are then fully specified by the list of reactive monomers carrying B molecules (*v*_1_), the list of pairs of monomers cross-linked (*v*_2_), and the Euclidean positions of the *N* monomers {*r*} = {**r**_1_, **r**_2_ *…* **r**_*N*_} (see Fig. 2). The partition function providing the statistical weight of each microstate reads as follows (if *β* = 1*/k*_*B*_*T* where *k*_*B*_ is the Boltzmann’s constant)

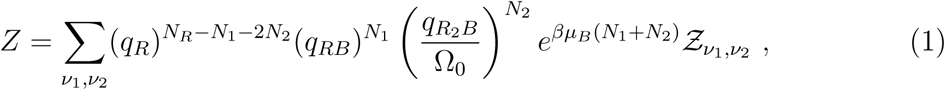

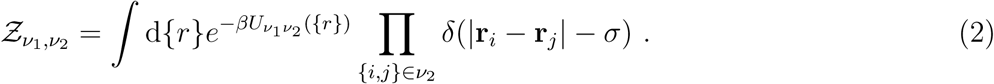

In Eq. 1, *q*_*X*_ (*X* = *R, RB*, and *R*_2_*B*, see Fig. 2a) are the internal partition functions of the reactive monomers (respectively, when free, carrying a linker, and cross-linked). At the same time, 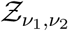 is the contribution accounting for the possible configurations of the chain backbone at a given *v*_1_, *v*_2_. A pair of monomers (*i* and *j*) cross–linked by a bridging molecule ({*i, j*} *∈ v*_2_) are constrained to stay at a fixed distance equal to *σ*. In Eq. 2, the integral over the chain’s backbone is constrained by the fixed bond condition, while 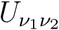 specifies the interactions between non-linked monomers as detailed below. Using the language of Ref. [30], Eqs. 1, 2 define a probability distribution on a stratification: a hierarchy of nested manifolds in 2(*N -* 1) *-N*_2_ + 3 dimensions (where 3 refers to the center of mass degrees of freedom).

As highlighted in a thread of investigations developing reaction ensemble Monte Carlo methods [34, 35], the internal partition functions are linked to the association equilibrium constant (*K*_*a*_) measured in diluted mixtures of B molecules and R monomers, the latter not constrained to stay on a chain (Fig. 2a). Specifically, the partition function of a single molecule of type *X* (*X* = *R, B, RB*, and *R*_2_*B*) is *V · q*_*X*_, where *V* is the volume of the system corresponding to the configurational space accessible to the center of mass of *X* treated as a classical variable. In diluted conditions, we can calculate the chemical potential (*µ*_*X*_) from the partition function of *N*_*X*_ molecules as follows

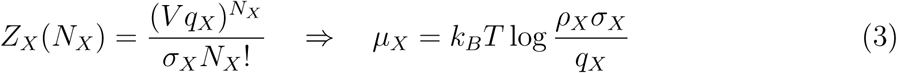

where *ρ*_*X*_ is the density of the molecule *X* and *σ*_*X*_ the symmetry order of the coarse-grained representation of *X* (*σ*_*R*_ = *σ*_*RB*_ = 1, 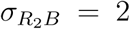, and *σ*_*B*_ = 1 as we treat B molecules implicitly). We now consider the two reactions leading to the formation of RB and R_2_B molecules: R+B ⇌ RB and RB+R ⇌ R_2_B. In equilibrium conditions, the sum of the chemical potentials of the educts should be equal to the same quantity calculated for the products: *µ*_*R*_ + *µ*_*B*_ = *µ*_*RB*_ and *µ*_*R*_ + *µ*_*RB*_ = *µ*_*R*2_*B*. When using Eq. 3, these equations allow expressing the internal partition functions in term of the association constant *K*_*a*_ as follows

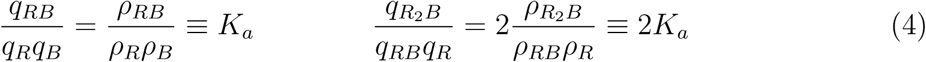

where we assume that the two terminals of the B molecules react with the same strength when binding the first and the second R monomer. When considering reactive monomers tethered to a chain, we can use Eqs. 4 in Eq. 1 to calculate the statistical weight 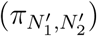 of binding a B molecule to the chain (*N*_1_ → *N*_1_ + 1) and of forming a loop (*N*_1_ → *N*_1_ *-* 1 and *N*_2_ → *N*_2_ + 1) relative to the one of a reference state 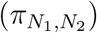

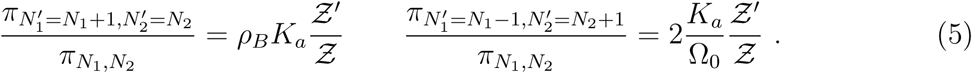

The previous expressions clarify how *ρ*_*B*_ and *K*_*a*_ are the only parameters required to parametrize the model. Instead, the configurational contributions *Ƶ′/Ƶ* are sampled using Monte Carlo simulations of the coarse-grained model, as detailed in the next section. Ω_0_ (see Eqs. 1, 5) is the configurational volume available to the orientational degree of freedom of an R_2_B dumbbell (Ω_0_ = 4*πσ*^2^). In Eq. 1, we divide 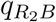 by Ω_0_, given that both *Z* (Eq. 2) and 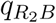 (Eq. 3) account for all possible orientations of an R_2_B molecule. Notice that Ω_0_ does not divide *q*_*RB*_ in Eq. 1 as Z does not depend on the orientation of the B molecules attached to the chain.

We now discuss the interactions between non–linked monomers 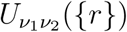 (Eq. 2). We decompose 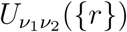 into the sum of pair potentials, 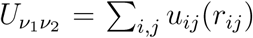, *r*_*i,j*_ = |**r**_*i*_ − **r**_*j*_| where *i* and *j* are not linked. We consider precursors in good solvent regimes. In particular, reactive and non-reactive monomers interact through purely repulsive potential *u*_*i,j*_ = *u*^*R*^. However, when considering interacting bridges (see Fig. 1c), interactions between reactive monomers bound to a B molecule (either belonging to an RB or R_2_B complex) feature an attractive well modeled using a cut and shifted Lennard–Jones potential, *u*_*i,j*_ = *u*^*A*^.

Specifically, we have

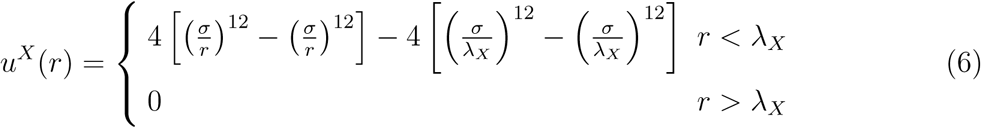

with *X* = *A, R* and *λ*_*R*_ = 2.5*σ*,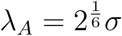.

### The simulation strategy

First, we consider the MC moves that change the connectivity state of the chain *v*_2_ by reacting complementary monomers. When attempting to form a new loop (for instance, by reacting monomers *i* with *j* in Fig. 3a), we start from deleting from the system a fraction of the chain (Γ) in the proximity of *j* (dashed segment in the top of Fig. 3a). We then generate a new segment configuration 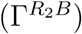 with *i* linked with *j* (Fig. 3a bottom). This process is done as proposed in [29] using methods growing chains with fixed endpoints [36–39]. In the reverse move, we delete a segment of the chain 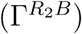 containing a randomly selected monomer forming a loop (*j*) and regrow a segment (Γ) without any loop. Acceptance rates are calculated as done in Configurational Bias Monte Carlo [40–42]. Importantly, we calculate the relative statistical weights of the two configurations of Fig. 3a using the second of Eqs. 5. The acceptance rates become then a function of the association constant *K*_*a*_. We refer to SM Sec. 2.3 for the details of the algorithm.

**Figure 3.**
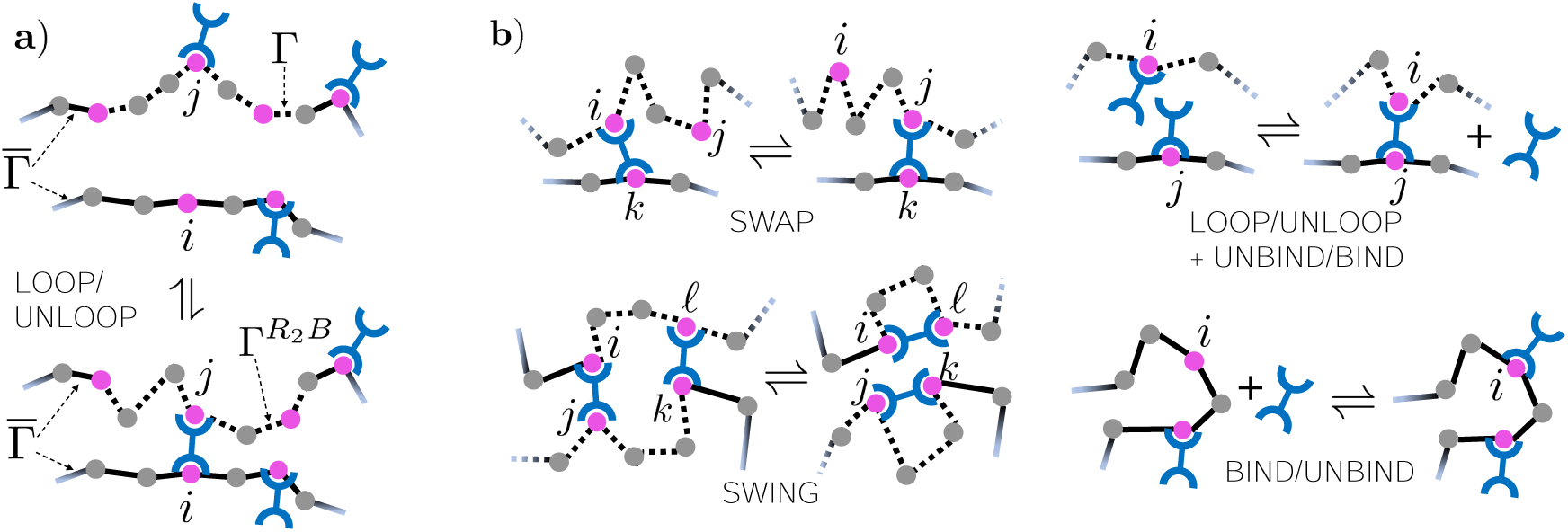
**a)** Loop/unloop MC moves react complementary monomers (*i* and *j*) while updating a segment of the chain encompassing *j* (highlighted using dotted line). **b)** MC moves involving multiple reaction events or changing the number of B molecules attached to the chain.

We also used MC moves that implement multiple binding/unbinding events simultaneously. In the Swap MC move (Fig. 3b), monomer *k* detaches from *i* and sequentially binds a second complementary monomer *j* [29]. In the Swing MC move (Fig. 3b), two pairs of reacted monomers exchange their partners [29]. In this work, we also use an MC move in which two reactive monomers carrying a B molecule react while simultaneously detaching one of the two bridges (loop/unloop+unbind/bind in Fig. 3b). B molecules are reversibly attached to the chain using the binding/unbinding moves (Fig. 3b). Contrary to the swing and the swap move, the binding/unbinding moves do not update the configuration of the backbone {*r*}. The details of the MC moves of Fig. 3b are reported in SM Sec. 2.2 (bind/unbind) and SM Secs. 2.4, 2.5, and 2.6 (respectively, for the loop/unloop+binding/unbinding, the swap, and the swing).

To further relax the system, we also employ standard MC moves that update the backbone of the polymer without affecting *v*_1_ and *v*_2_. Specifically, we use pivot and double pivot moves that rotate fractions of the chains. Segments of the chain are also regrowth using the CBMC method (as used in the loop/unloop move) without changing the connectivity state of the system. A dedicated MC move displaces, and reorients reacted complexes (R_2_B) while regrowing a fraction of the surrounding network. In each MC cycle, we randomly perform one of the previous moves.

### Comparison with existing methodologies

Simulation schemes to study the statistical properties of chains carrying reactive units are finding applications in nanotechnology, for instance, to self-assemble polymeric nanoparticles [43]. Molecular Dynamics simulations have been used to study the morphological properties of the nanoparticles with an emphasis on finding protocols leading to maximal chain compaction [43, 44]. In most of these studies, cross-linking between complexes is irreversible. Refs. [26, 45] introduced a three-body potential that allows implementing swap-like moves in Molecular Dynamics simulations. Recently, Ref. [30] introduced a general MC scheme which allows adding/removing holonomic constraints reversibly. Inter-molecular, reversible linkers are also used to enforce topological entanglement when studying polymer melts using soft potentials [46]. In this respect, we note that the current version of our simulation method, the chains are crossable. On the one hand, the crossability of the chromatin fiber is guaranteed by the action of dedicated enzymes. Secondly, topological constraints could be included in our methodology using linking numbers.

Chains carrying monomers featuring selective interactions are currently used to study condensation of multivalent proteins into membraneless bodies [22–24]. These contributions study the molecular determinants underlying the aggregation of multivalent proteins comprising short folded domains linked by intrinsically disordered regions. In particular, the LASSI package [24] employs lattice models in which folded domains are mapped into beads forming reversible linkages while disordered regions into strings of beads interacting through non-specific interactions. The lattice model is parametrized by atomistic simulations and is sequence-dependent [47]. Relevant to the findings of this work, non-specific interactions between linkers could lead to microphase separation in multicomponent systems [23] and affect the physical mechanism underlying chain aggregation [22]. The MC moves presented in the previous section could be readily adapted to the LASSI setting. Finally, simulations based on field-theoretic methods [48] have also been used to study supramolecular polymer physics [25].

## RESULTS AND DISCUSSION

### Non-Interacting Bridging Factors (NIBs): Monofunctional Chains

In this section, we quantify how the bridges’ density (*ρ*_*B*_) and the association strength (*K*_*a*_) regulate the number of reactive monomers carrying linkers (*N*_1_) as well as the number of loops (*N*_2_). We then correlate *N*_1_ and *N*_2_ with the morphological properties of the chains, namely the gyration radius, the distribution of the loop length, and the averaged connectivity map.

When forbidding the formation of loops, the bindings of bridges to different reactive monomers become independent events, and the probability that a reactive monomer carries a B molecule becomes equal to (see the discussion after Eq. 8)

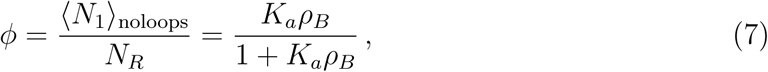

where ⟨ *…* ⟩_noloops_ denotes a constrained average with *N*_2_ = 0 as obtained using Eq. 1. Motivated by this observation, in the following, we use the variable *ϕ* to discuss our results. Fig. 4a and SM Fig. S1a study the fraction of reactive monomers carrying a linker without forming a loop (⟨*N*_1_⟩*/N*_*R*_) as a function of *ϕ* when changing *K*_*a*_ (Fig. 4a) and *f* (SM Fig. S1a). As expected, ⟨*N*_1_⟩ increases with *ϕ* from *N*_1_ = 0 (for *ϕ* = 0) to *N*_1_ = *N*_*R*_ (for *ϕ* = 1). When increasing the association constant, ⟨*N*_1_⟩ starts to deviate from the linear behavior predicted by Eq. 7 due to loop formation. In particular, at a given *ϕ*, loops become more favorable at high values of *K*_*a*_ given that the formation of loops at a given *N*_1_ + *N*_2_ is only controlled by the association constant and free-energy terms (discussed below) which do not depend on *ρ*_*B*_ (see Eq. 1). In Fig. 4b and SM Fig. S1b, we study the number of loops as a function of *ϕ*. As anticipated above, ⟨*N*_2_⟩ increases with *K*_*a*_ (Fig. 6b) and the degree of functionalization *f* (if *f* ≤ 0.5, see SM Fig. S1b). Importantly, the number of loops is non-monotonic in *ϕ* and goes to zero when *ϕ* tends towards *ϕ* = 1. This observation underlies the unfolding of the chain when overexpressing bridging molecules. Entropic terms regulate the opening of a loop in favors of two R monomers carrying two bridging molecules. The process is purely entropic as the number of reacted monomers (and therefore the number of *K*_*a*_ terms entering the Boltzmann distribution, see Eq. 1) in the two competing microstates does not change. For high values of *ρ*_*B*_, the system attempts to maximize the number of bridging molecules present on the chain, therefore opening loops.

**Figure 4.**
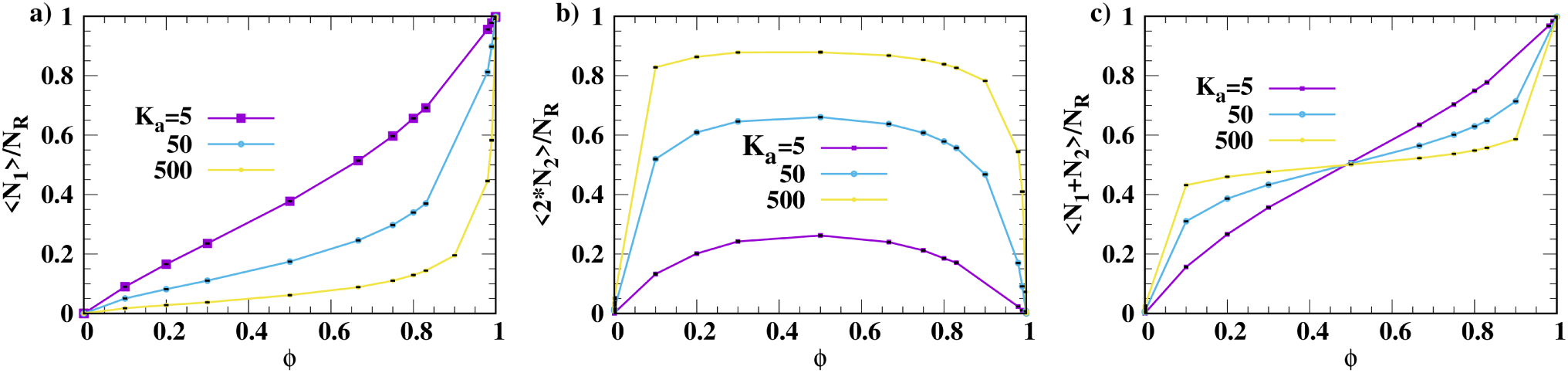
**a**-**c)** Fraction of reactive monomers carrying a linker, forming a loop, and number of B molecules attached to the chain *per* R monomer as a function of *ϕ. N* = 1000 and *f* = 0.5. In all cases, the errorbar is smaller than the size of the symbols.

**Figure 5.**
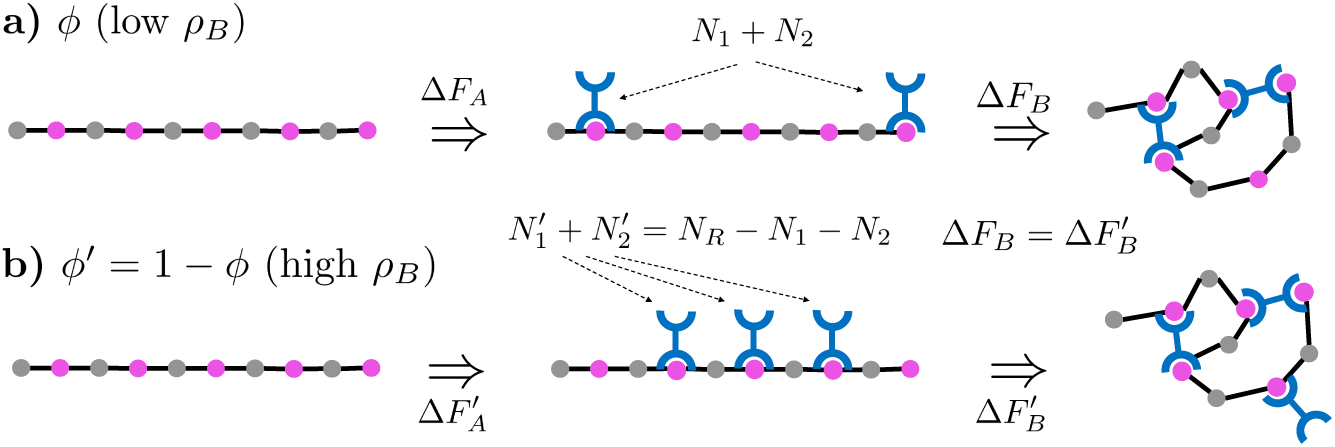
A two-step pathway to calculate the free energy of the system with a given *N*_1_ and *N*_2_. Δ*F*_*A*_ is the free energy of attaching *N*_1_ + *N*_2_ B molecules to the fiber, while Δ*F*_*B*_ accounts for the loops’ contribution to the free energy.

**Figure 6.**
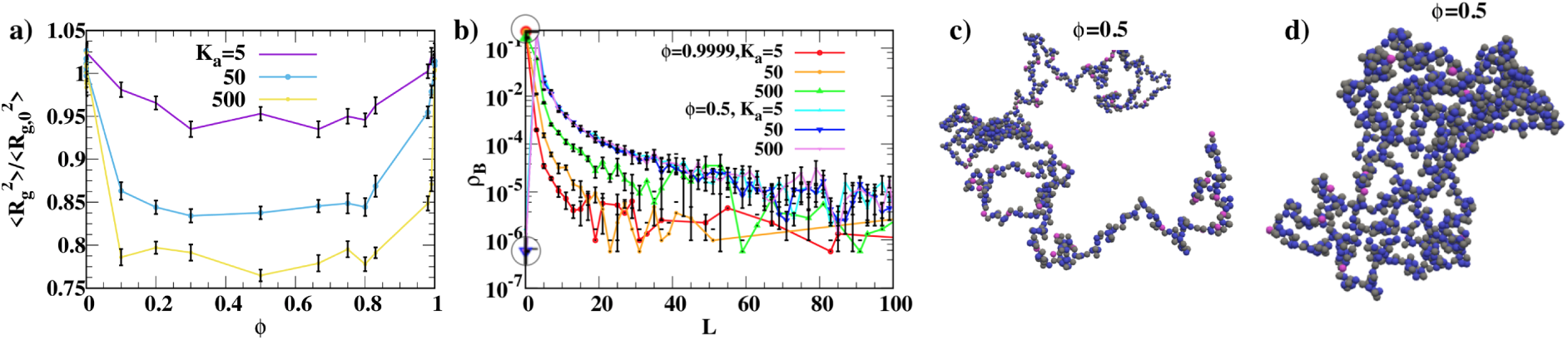
**a)** Averaged radius of gyration 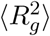 of chains folded by NIBMs as a function of *ϕ*. 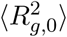 is the averaged gyration radius in a system without B molecules. **b)** Loop length distribution for different values of *K*_*a*_ and *ϕ*. In **a)** and **b)**, we use *N* = 1000, *f* = 0.5, and calculate errorbars using 50 independent simulations consisting of 5 *·* 10^5^ MC cycles. **c**) and **d**) are snapshots with *K*_*a*_ = 500, when, respectively, using and not using swap and swing MC moves (Fig. 3b). Grey, pink, and blue beads represent, respectively, non reactive monomers, not–looped R monomers, and looped monomers.

Intriguingly, for all values of *K*_*a*_ and *f* reported in Fig. 4b and SM Fig. S1b, the plots of ⟨*N*_2_⟩ as a function of *ϕ* are symmetric with respect to the axis *ϕ* = 1*/*2. To understand this result and make new predictions, in Fig. 5, we present a pathway to estimate the free energy of two systems at different *ρ*_*B*_ (corresponding to a given *ϕ* and *ϕ′* = 1 *-ϕ* with *ϕ′ > ϕ*) and same *K*_*a*_. Taking as a reference state a polymer without any B molecule, we decompose the free energy of the system into the contribution of putting *N*_1_ + *N*_2_ B molecules on the chain (Δ*F*_*A*_) and the one arising from making *N*_2_ loops (Δ*F*_*B*_) without attaching any extra linker to the polymer. Δ*F*_*A*_ can be calculated as

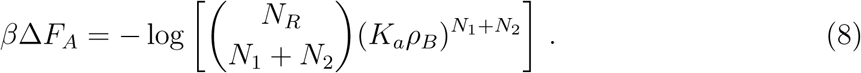

In Eq. 8, the binomial term counts the ways of distributing *N*_1_ + *N*_2_ bridges within *N*_*R*_ R monomers, and *K*_*aρB*_ is the statistical weight of each microstate as obtained using Eq. 1 and Eqs. 4. (Notice that Eq. 7 follows from Eq. 8.) Using Stirling’s approximation, we obtain

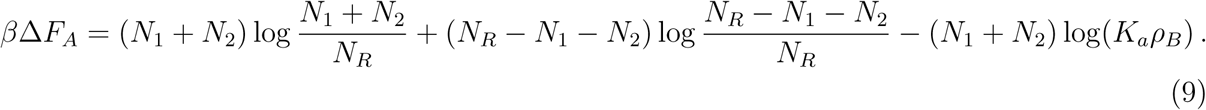

The calculation of Δ*F*_*B*_ is not feasible. Mean-field estimates of Δ*F*_*B*_ are also tricky because, in contrast to systems with ligand–presenting colloids, reactions between complementary monomers can hardly be treated as independent events. Even though analytic expressions of this term are not available, we can prove that Δ*F*_*B*_ is the same for the two systems considered in Fig. 5. This claim follows from the fact that, in the two systems, the numbers of R, empty monomers and R monomers carrying a bridge are exchanged. It follows that the combinatorial and configurational terms entering the calculation of Δ*F*_*B*_ are the same in the two cases, as B molecules do not affect the local morphology of the chain. Therefore, in general, we can write the following equality

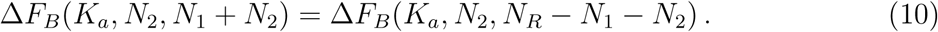

The previous equation explains the reason why ⟨*N*_2_⟩ is symmetric with respect to the axis *ϕ* = 1*/*2: At a given value of *N*_1_ + *N*_2_, the two terms of Eq. 10 are minimized by the same value of *N*_2_.

We can push our analysis a step further and predict that the total number of linkers on the chain (⟨*N*_1_ + *N*_2_⟩) is an antisymmetric function of *ϕ*. The most likely number of *N*_1_ and *N*_2_ follows from saddle point equations

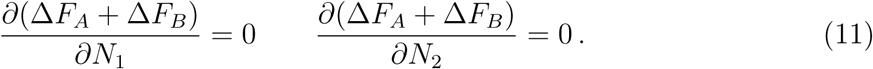

When developing the first of these equations using Eq. 8, we obtain

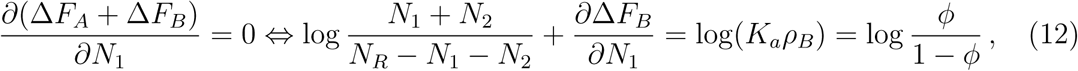

where we use Eq. 7 to express *K*_*aρB*_ in terms of *ϕ*. Under the transformations *N*_1_ + *N*_2_ → *N*_*R*_ *-N*_1_ *-N*_2_ and *ϕ* → 1*-ϕ*, all the terms of the second equation change sign. It follows that ⟨*N*_1_ + *N*_2_⟩ is an antisymmetric function of *ϕ* along the axes *ϕ* = 1*/*2 and *N*_1_ + *N*_2_ = *N*_*R*_*/*2. We verify this prediction in Figs. 4c, SM Fig. S1c using simulation results. In particular, the total number of B molecules on the chain increases with *ϕ*. Similar to Fig. 4c and SM Fig. S1c, when increasing the values of *K*_*a*_ and *f*, the plots deviate from a linear behavior (corresponding to *N*_2_ = 0). For *ϕ <* 1*/*2, ⟨*N*_1_+*N*_2_⟩ *> ϕ* as two R monomers can cooperatively stabilize a B molecule through the formation of a loop. Instead, for *ϕ >* 1*/*2, ⟨*N*_1_ + *N*_2_⟩ *< ϕ* as multiple B molecules are shared by pairs of R monomers. When *ϕ* → 0 and *ϕ* → 1, the behavior of ⟨*N*_1_ + *N*_2_⟩ is dominated, respectively, by ⟨*N*_2_⟩ and *N*_*R*_ *-* ⟨*N*_2_⟩.

Importantly the previous arguments rely on the fact that we did not consider non-specific interactions between linker molecules and the chain’s backbone. For instance, in the presence of steric interactions, Eq. 10 (and similarly Eq. 8) would not be valid as more bridges on the chain would increase the configurational cost of folding chains. Although we expect the general trend to remain unaffected by non-specific interactions, the simulation methodology can easily be adapted to account for these terms.

In Fig. 6a, we study the averaged squared radius of gyration 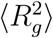 as a function of *ϕ* for *f* = 0.5. We observe that 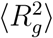 is non-monotonic in *ϕ* with the chains that reswell when the system tends towards *ϕ* = 1. This behavior mirrors the trends observed for the number of bridges (see Fig. 4b) and clarifies how intra–molecular linkages drive the compaction of the fiber. In particular, 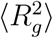 decreases when increasing the value of *K*_*a*_ as more loops become stable. In Fig. 6b, we report the probability of forming a loop made by L segments for different values of *K*_*a*_ and *ϕ*. The *L* = 0 value refers to the probability for an R monomer to be unlooped. Consistently with Fig.6a, increasing *K*_*a*_ decreases the probability of finding monomers unpaired. Moreover, the system attempts to minimize the length of loops as a result of the higher configurational cost of forming longer loops. As a result, the loop length distribution sharply decreases with *L*. Longer loops are expected when reducing the non–specific repulsion between monomers (*u*^*R*^ in Eq. 6). However, the system can feature persistent longer loops unless employing MC moves that change the length of a loop in a single step [29]. This result is shown in Fig. 6c and 6d, where we report two fiber structures when using (Fig. 6c) or not (Fig. 6d) the swap and the swing MC move (Fig. 3b). In the second case, we obtain a much more compact structure with longer loops that persist during the simulation. Longer loops will also occur when using semiflexible backbones. However, we stress how the unfolding of the fiber at high *ρ*_*B*_ is not model dependent and will be found in all systems featuring non–interacting bridging molecules. Consistently with the fact that thermodynamic states only feature short loops, SM Figs. S2, S3 show how the results of Figs. 4, 6a are not affected by the chain’s length.

### Non-Interacting Bridging Factors (NIBs): Bifunctional Chains

In this section, we consider chains carrying two types of R monomers (R_1_ and R_2_) reacting with a single type of B molecules with different affinities (defined as 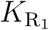 and 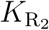). We aim to study if some degree of dispersity in the specific interactions between R monomers and B molecules could broaden the range of *ρ*_*B*_ corresponding to compact chains. We consider chains made of repeated blocks in which half of the monomers are reactive. Each block comprises one monomer of type R_1_ and four of type R_2_, interposed with inert monomers (see inset of Fig. 7b). In Fig. 7b, we report the fraction of reactive monomers forming a bridge as a function of *ρ*_*B*_ while keeping the association constant of the strongest monomer (*R*_1_) equal to 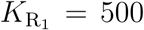. We change the association constant between B and *R*_2_ 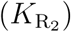 from 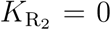 to 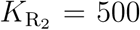. The two limiting cases (dashed lines in Fig. 7b) correspond to monofunctional systems with, respectively, *f* = 1*/*10 (R_2_ monomers stop being reactive) and 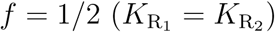.

**Figure 7.**
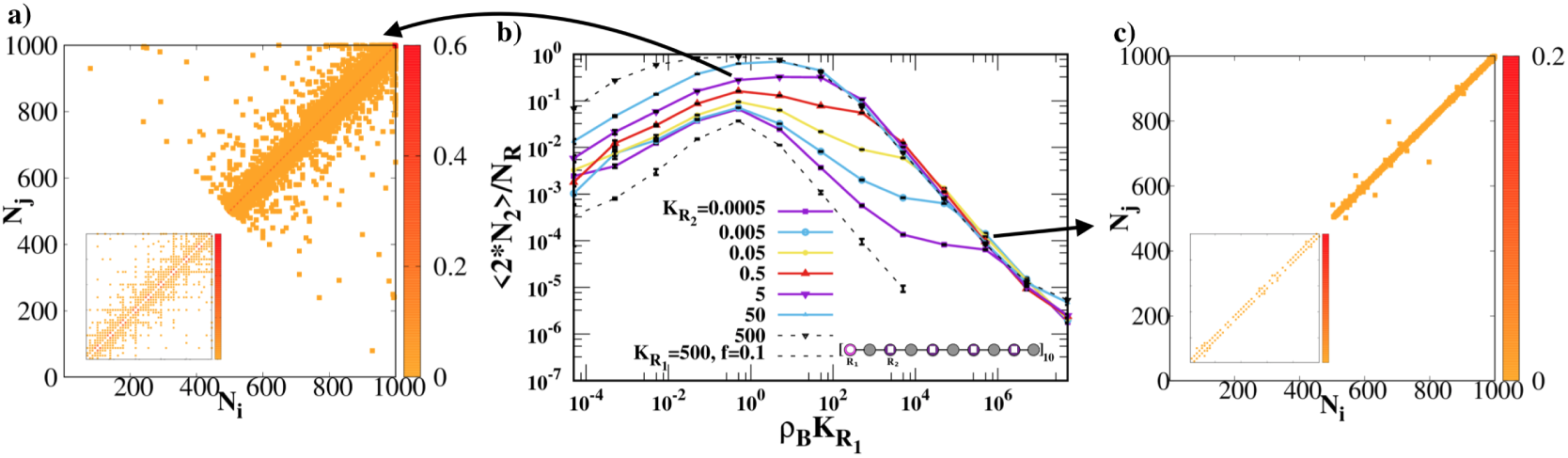
**a)** and **c)** Averaged connectivity maps of bifunctional chains with *f* = 0.5 and 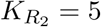. In **a)**, 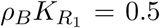, while in **c)** 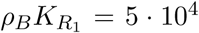. The inset shows the connectivity map of the segment made of the first 100 beads. **b)** Fraction of reactive beads forming a loop as a function of _*ρB*_ at different 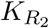 for 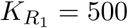.

We recover the same phenomenology found in the monofunctional case. In particular, intra-molecular bridges disappear, both for large and small values of *ρ*_*B*_. An inspection of the connectivity maps (see Fig. 7a) shows how, for intermediate values of *ρ*_*B*_, the chains predom-inantly form loops between *R*_1_ and *R*_2_ monomers. Reacting *R*_1_ with *R*_2_ allows minimizing the configurational costs of forming long loops while prioritizing the binding of B molecules to R_2_ monomers. When increasing *ρ*_*B*_, the system attempts to maximize the number of B molecules on the chains. In these conditions, the connectivity maps (see Fig. 7c) do not feature the texture typical of copolymer architectures (like in Fig. 7a). In particular, the few short loops involve with equal probability *R*_1_ and *R*_2_. Interesting, all curves (with 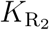 > 0) collapse onto the monofunctional case (with *f* = 0.5) in the large *ρ*_*B*_ limit. This general finding clarifies how purely entropic terms control the unfolding transition. However, the unfolding transition is not entropic in the case of bridging molecules formed by dimers (like in the case of associating YY1 [49] or CTFC binding factors [50]). Indeed, in the latter case, two passivated R monomers would carry two pairs of dimers as compared to a single dimer entering a loop. Therefore, we predict how large dimerization constants would accelerate the unfolding transition.

### Interacting Bridging Molecules (IBMs)

In this section, we consider the case in which B molecules interact as due, for instance, to multivalent interactions [16]. Here, NIBMs can condensate around multiple R sites that can therefore cluster. Following the strategy of Ref. [9], we drive the clustering of R monomers carrying B molecules using the LJ potentials introduced in Eq. 6.

Fig. 8a reports the averaged gyration radius of the IBM model as a function of *ϕ* for different degrees of functionalization *f*. As compared to Fig. 6, 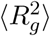 rapidly decreases with *ϕ* and then exhibits a plateau for *ϕ ⪆* 0.1. The sharp decrease of 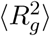 in *ϕ* is due to a cooperative effect in which B molecules are colocalized by R monomers and stabilized by their mutual interaction. We verify this claim in SM Fig. S4, where we compare the fraction of R monomers carrying a B molecule in the NIBM and the IBM model. More B molecules are found on the chain in the case of the IBM since, in this model, R monomers can simultaneously interact with multiple partners. This observation also explains the fact that chains are more compact in the NIBM model (compare Fig. 8a with Fig. 6a). Similar findings have been reported in synthetic systems, where scholars struggled to self-assemble compact polymeric nanoparticles using intra-molecular bridges [43].

**Figure 8.**
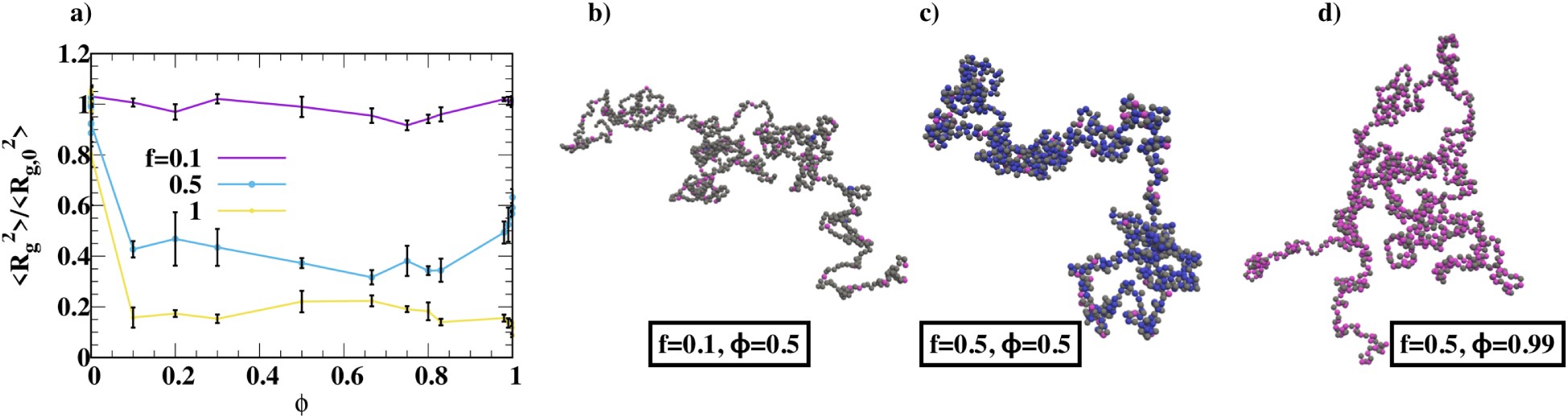
**a)** Averaged radius of gyration 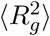 of chains folded by IBMs as a function of *ϕ* for *K*_*a*_ = 500. 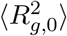 is the averaged gyration radius in a system without bridging molecules. We have calculated error bars using 50 independent simulations consisting of 5*·*10^5^ MC cycles. **b)**-**d)** Typical snapshots for *K*_*a*_ = 500 and different *f, ϕ*. Grey, pink and blue beads represent, respectively, non reactive monomers, not–looped R monomers, and looped monomers.

Fig. 8a does not exhibit the unfolding transition, and the chains stay compact even for *ϕ* = 1. This result follows from the fact that monomers carrying B molecules are not passivated but can still interact as a result of the interactions between B molecules. This result is confirmed by snapshots showing how for *f* = 0.5 and different values of *ϕ*, the morphology of the fiber is comparable (see Fig. 8c and 8d and SM Fig. S5 for the same configurations in which we only report beads bound to a B molecule). On the other hand, smaller values of *f* increase the size of the chain as R monomers interact with fewer partners (see Fig. 8a and 8b).

## CONCLUSION

We presented a chromatin model consisting of a chain carrying reactive monomers folded by bridging (B) molecules dispersed in solution (at a fixed density, *ρ*_*B*_) reversibly attaching the chromatin fiber. We studied the effect of *ρ*_*B*_ and the association constant between B molecules and the reactive monomers on the morphological properties of the chain. For intermediate values of *ρ*_*B*_, the chromatin folds due to the formation of loops comprising B molecules cross-linking two reactive monomers. Instead, overexpression of bridging factors can lead to the unfolding of the fiber. We highlighted the generality of this unfolding transition using thermodynamic considerations. Importantly, the unfolding transition is peculiar to non-interacting, bridging molecules. Instead, for models featuring B molecules that tempt to phase separate when colocalized on the fiber, chains are much more compact (even for high values of *ρ*_*B*_) as a result of the fact that interactions between reactive monomers are not valence-limited.

Our results are supported by Monte Carlo methodologies, which, as compared to Molecular Dynamics simulations, allow reconfiguring the topology of the networks through dedicated moves, therefore enabling efficient sampling between configurations featuring different connectivity states. Equilibrium configurations feature almost exclusively short loops. Short loops are thermodynamically stable because they minimize excluded volume interactions and configurational costs. However, we noticed that it is somehow difficult to relax the system towards the ground state. In particular, longer loops (resulting in much more compact chains) will persist unless using dedicated MC moves implementing multiple reactions in a single step.

The results presented in this paper allow inferring the microscopic mechanisms driving compaction from experimental results.

## Supporting information

Supplemental figures and simulation details

## AUTHOR CONTRIBUTIONS

BMM, IM and BO designed the research. BO and IM developed the program. IM carried out all simulations, analyzed the data. IM and BMM wrote the article.

## ACKNOWLEDGMENTS

We thank K. Rippe and F. Erdel for pointing our attention towards Ref. [17] and for suggesting the study of Fig. 7. Financial support was provided by an A.R.C grant of the *Fédération Wallonie-Bruxelles*. Computational resources have been provided by the Consortium des Equipements de Calcul Intensif (CECI), funded by F.R.S.-FNRS under Grant No. 2.5020.11. The authors declare no conflict of interest.

