## Supplemental figures and simulation details for "Unfolding of the chromatin fiber driven by overexpression of bridging factors"

### 1 Supplementary Figures (Figs. S1-S5)

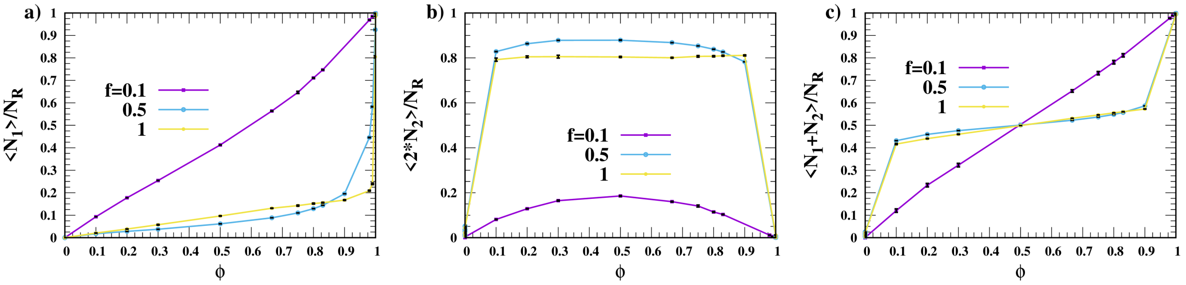


Figure S1: **a-c)** Fraction of reactive monomers carrying a linker, forming a loop, and number of bound B molecules *per* R monomer at $K_a = 500$ as a function of $\phi$ for different degrees of functionalization (see legends). In all cases, the error is smaller than the size of the symbols.

### 2 Details of the Simulation Method

In Sec. 2.1, we explain the algorithm to generate fractions of polymers with fixed ends. We then describe the MC moves of Main Fig. 3, namely, the bind/unbind move (Sec. 2.2), the

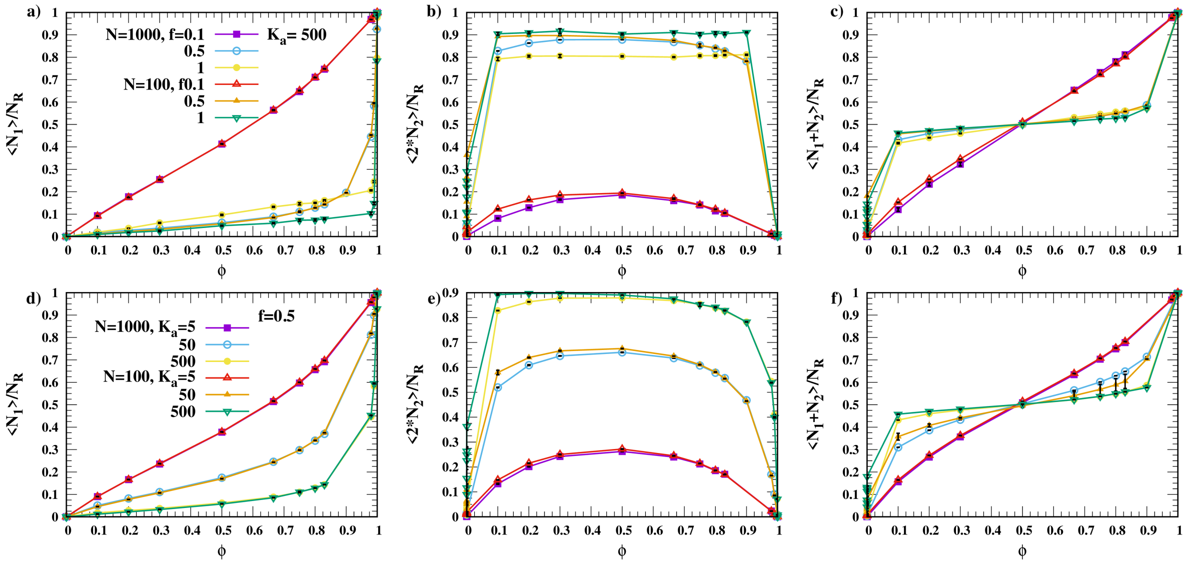


Figure S2: **a-c)** Same as in Fig. S1a-c, also including chains with $N = 100$ monomers. **d-f)** Same as in Main Fig. 4a-c, also including chains with $N = 100$ monomers.

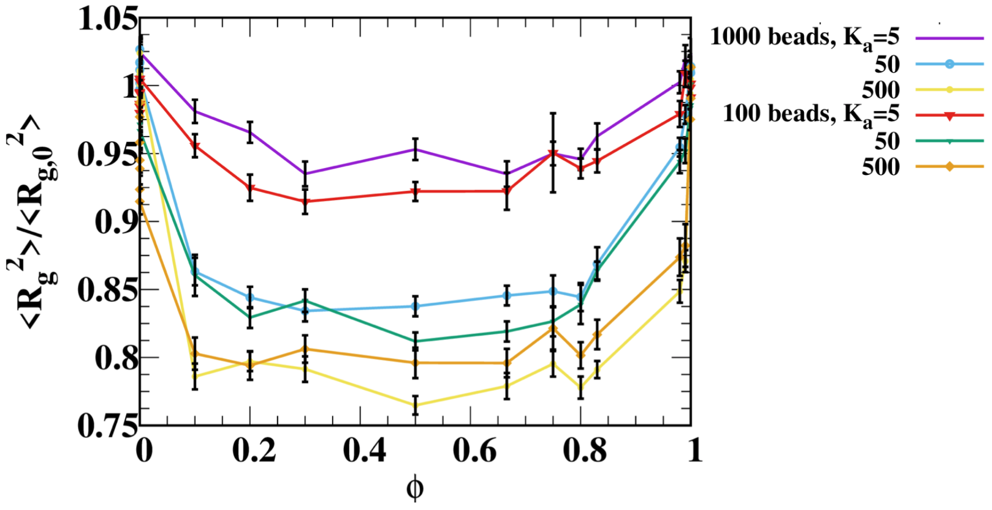


Figure S3: Same as in Main Fig. 6a, also including chains with $N = 100$ monomers.

loop/unloop move (Sec. 2.3), the loop/unloop+unbinding/binding move (Sec. 2.4), the swap move (Sec. 2.5), and the swing move (Sec. 2.6).

### 2.1 Sampling chains with fixed ends

In Fig. S6, we explain how we generate a new section of the chain made of $N_c$ monomers, $\Gamma^{I,F}_{\text{new}} = \{\mathbf{r}_{m_1}, ..., \mathbf{r}_{m_{N_c}}\}$, between two fixed terminals $\mathbf{r}_I$ and $\mathbf{r}_F$ ($I = m_1 - 1$ and $F = m_{N_c} + 1$).

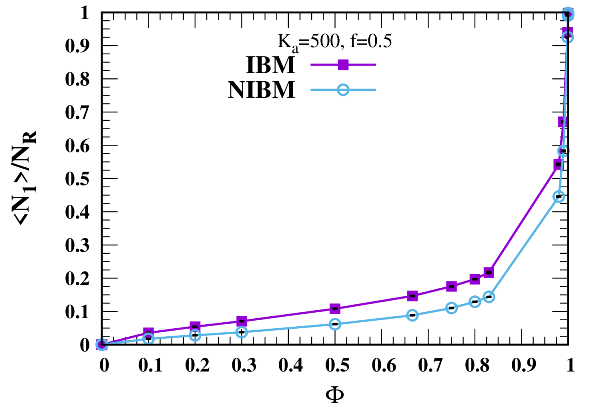


Figure S4: Fraction of reactive monomers carrying a B molecule for $f = 0.5$ and $K_a = 500$ as a function of $\phi$ for chains folded by NIBMs and IBMs.

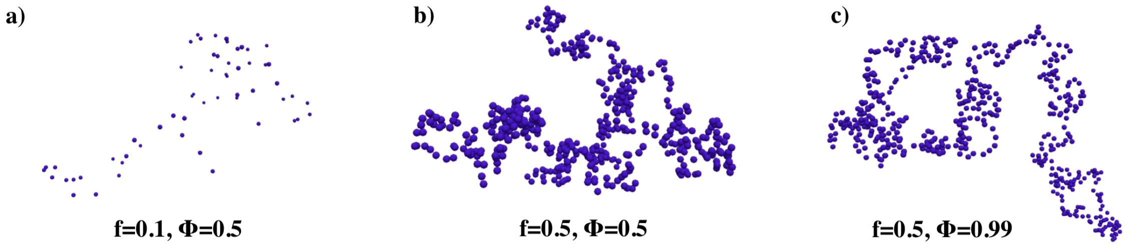


Figure S5: **a**-**c**) Beads bound to a B molecule for the configurations of Main Fig. 8b-d.

$m_1$ and $m_{N_c}$ are, respectively, the first and the last monomer of segment $\Gamma^{I,F}_{\text{new}}$. The beads are inserted sequentially by using the Rosenbluth method [7, 5, 6], starting from $m_1$ ($m_1 = I+1$). The growth of the chain is biased towards the final terminal $F$ by guiding functions $p$. In this work, we employ the end-to-end distance distributions of Freely-Jointed Ideal Chains (FJICs) as guiding functions [9, 11].

Fig. S6 explains the growing procedure. The insertion of monomer $m_i$ follows the standard method used in Rosenbluth sampling. We define the bond segment $\mathbf{u}_i$ as $\mathbf{u}_i = \mathbf{r}_{m_i} - \mathbf{r}_{m_{i-1}}$ and generate $k$ trial bond segments $\mathbf{u}_i^n$ (with $n = 1, ..., k$) using $p^i_{\text{trial}}$ defined as follows

$$p^i_{\text{trial}} = \begin{cases} \frac{p(N_c-i+1,|\mathbf{r}_{m_{i-1},F}+\mathbf{u}_i^n|)}{\Omega_0 p(N_c-i+2,|\mathbf{r}_{m_{i-1},F}|)} & \text{if } i < N_c \\ \frac{1}{2\pi} & \text{if } i = N_c \end{cases} \tag{1}$$

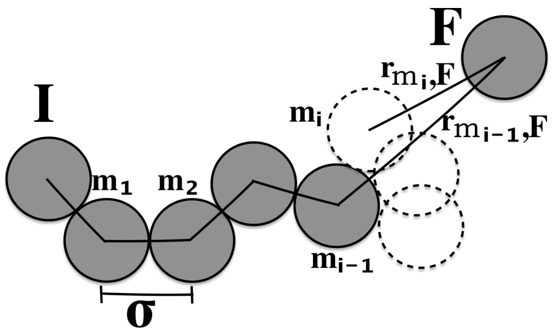


Figure S6: Growing procedure of a chain section $\Gamma^{I,F}_{\text{new}}$ made of $N_c$ monomers between monomer $I$ and $F$. The distance between the trial monomers $m_i$ (dashed spheres) and the end monomer ($|\mathbf{r}_{m_i,F}|$), and the number of monomers not yet inserted ($N_c - i + 1$) are used to bias the growth of the chain section towards $F$.

where $p(M, r)$ is the end-to-end distance ($r$) distribution of an FJIC made of $M$ monomers. The normalization of $p^i_{trial}$ follows from the following identity (Markov property)

$$\int \mathrm{d}\mathbf{u} \cdot p(N, |\mathbf{r} + \mathbf{u}|) = \Omega_0 \cdot p(N + 1, |\mathbf{r}|), \tag{2}$$

where $\Omega_0 = 4\pi\sigma^2$, and the integral of $\mathbf{u}$ is taken over a sphere of diameter $\sigma$ centered in (0, 0, 0). Monomer $N_c$ is inserted through a crankshaft move [1, 3] in which we sample the azimuthal angle $\phi$ of $\mathbf{r}_{m_{N_c}}$ pivoting around the axis $\mathbf{r}_{F,m_{N_C-1}}$ ($\mathbf{r}_{F,m_{N_C-1}} = \mathbf{r}_F - \mathbf{r}_{m_{N_c-1}}$).

In the next step, we select $\mathbf{u}_i$ within the $k$ trial bond segments ($\mathbf{u}_i \in \{\mathbf{u}^n_i\}_{n=1,\cdots k}$) with a probability $p^i_{\text{select}}(\mathbf{u}^n_i)$

$$p^i_{\text{select}} = \begin{cases} \frac{\exp[-\beta\, u_{m_i}]}{W^{(i)}_{u_i}} & \text{if } i < N_c \\ \frac{\exp[-\beta\, u_{m_{N_c}}]}{\Lambda^{(N_c)}_{u_{N_c}}} & \text{if } i = N_c \end{cases} \tag{3}$$

The Rosenbluth factors, $\Lambda^{(N_c)}_{\mathbf{u}_{N_c}}$ and $W^{(i)}_{\mathbf{u}_i}$, normalize $p^i_{\text{select}}$ and read as follows [8]

$$W^{(i)}_{\mathbf{u}_i} = \sum_{n=1}^{k} \exp[-\beta u^n_{m_i}] \qquad \Lambda^{(N_c)}_{\mathbf{u}_{N_c}} = \sum_{n=1}^{k} \exp[-\beta u^n_{m_{N_c}}] \tag{4}$$

In Eqs. 3, 4, $u^n_{m_i}$ denotes the non-specific interaction of the trial monomer $\mathbf{r}^n_{m_i} = \mathbf{r}_{m_{i-1}} + \mathbf{u}^n_i$ (calculated using Main Eq. 6) with the rest of the chain, including the fraction of $\Gamma^{I,F}_{\text{new}}$ already grown. In particular

$$u^n_{m_i} = \begin{cases} \sum_{j=1}^{\min[I,m_i-2]} u_{m_i,j}(r_{m_i,j}) + \sum_{j=\max[F,m_i+2]}^{N} u_{m_i,j}(r_{m_i,j}) & \text{if } i < 3 \\ \sum_{j=1}^{\min[I,m_i-2]} u_{m_i,j}(r_{m_i,j}) + \sum_{j=\max[F,m_i+2]}^{N} u_{m_i,j}(r_{m_i,j}) & \\ \quad + \sum_{j=1}^{j<i-1} u_{m_i,m_j}(r_{m_i,m_j}) & \text{if } i \geq 3 \end{cases} \tag{5}$$

where $u_{i,j} = u^R$ or $u_{i,j} = u^A$ depending on the type of model (see Main Eq. 6). $u_{m_i}$ is the non-specific interaction of the selected segment.

Using $p^i_{\text{trial}}$ and $p^i_{\text{select}}$ (Eqs. 1, 3), we write the probability of generating a segment $\Gamma^{I,F}_{\text{new}}$ as follows

$$\begin{aligned} P_{\text{grow}}(\Gamma^{I,F}_{\text{new}}) &= \frac{1}{J(\Gamma^{I,F}_{\text{new}})} \prod_{i=1}^{N_c} p^i_{\text{trial}} \cdot p^i_{\text{select}} \\ &= \frac{\exp[-\beta\, U_{\Gamma^{I,F}_{\text{new}}}] p(2, r_{m_{N_c-1},F})}{2\pi J(\Gamma^{I,F}_{\text{new}}) \Omega_0^{N_c-1} W_{\Gamma^{I,F}_{\text{new}}} p(N_c+1, r_{I,F})} \\ W_{\Gamma^{I,F}_{\text{new}}} &= \Lambda^{(N_c)}_{\mathbf{u}_{N_c}} \prod_{i=1}^{N_c-1} W^{(i)}_{\mathbf{u}_i}\,, \end{aligned} \tag{6}$$

where $U_{\Gamma^{I,F}_{\text{new}}}$ is the configurational energy of $\Gamma^{I,F}_{\text{new}}$

$$\beta\, U_{\Gamma^{I,F}_{\text{new}}} = \sum_{i=1}^{N_c} u^n_{m_i}\,. \tag{7}$$

In Eq. 6, we considered a Jacobian term $J$ given that monomer $m_{N_c}$ is inserted by sampling the azimuthal variable $\phi$ at fixed $\mathbf{r}_{F,m_{N_c-1}}$. $J$ is then the Jacobian resulting from the change of coordinates from $\{\mathbf{r}_{m_{N_c-1}}, \phi\}$ to $\{\mathbf{u}_{N_c} \equiv \mathbf{r}_{m_{N_c}} - \mathbf{r}_{m_{N_c}-1}, \mathbf{u}_F \equiv \mathbf{r}_F - \mathbf{r}_{m_{N_c}}\}$ [1, 10, 2]. In particular, $J(\Gamma^{I,F}_{\text{new}}) = |\partial(\mathbf{u}_{N_c},\, \mathbf{u}_F)/\partial(\mathbf{r}_{m_{N_c-1},F},\, \phi)| = \sigma^2/r_{m_{N_c-1},F}$.

In the following sections, we will need to estimate the probability of generating an existing segment (labeled with $\Gamma^{I,F}_{\text{old}}$). To do that, we regrow $\Gamma^{I,F}_{\text{old}}$ by following the same procedure employed to generate $\Gamma^{I,F}_{\text{new}}$, except that the $k-th$ trial bond is taken equal to the position of the actual monomer $u_{m_i}$, which is also selected. The expression of $P_{\text{grow}}(\Gamma^{I,F}_{\text{new}})$ then follows from Eq. 6.

We will also need to grow segments of the chain ($\Gamma^I$) without fixed end monomer (see, for instance, $\Gamma_2$ in Fig. S7c-d). In this case, $p^i_{\text{trial}}$ is not biased by the guiding function $p(N, r)$. Instead, the $k$ trials are generated uniformly while the trials are sorted using the first of Eq. 3. $P_{\text{grow}}(\Gamma^I_{\text{new}})$ is then given by

$$\begin{aligned} P_{\text{grow}}(\Gamma^I_{\text{new}}) &= \frac{\exp[-\beta U_{\Gamma^I_{\text{new}}}]}{\Omega_0^{N_c} W_{\Gamma^I_{\text{new}}}} \\ W_{\Gamma^I_{\text{new}}} &= \prod_{i=1}^{N_c} W^{(i)}_{\mathbf{u}_i} \end{aligned} \tag{8}$$

where $U_{\Gamma^I_{\text{new}}}$ is the configurational energy of $\Gamma^i_{\text{new}}$. The probability of regrowing an old configuration $\Gamma^I_{\text{old}}$ is then $P_{\text{grow}}(\Gamma^I_{\text{old}})$. The regrowing procedure is analogous to what done in the case of segments with fixed ends.

### 2.2 Bind/Unbind MC move

In this section, we describe the MC move that, with equal probability, attempts either to bind or to detach a B molecule from the chain. When trying to attach a B molecule to the chain, we select one within the $N_R - N_1 - 2N_2$ free reactive monomers with uniform probability

$$K(N_1 \to N_1 + 1) = \frac{1}{N_R - N_1 - 2N_2}\,. \tag{9}$$

Similarly, when attempting to detach a B molecule from the chain, we select one within the $N_1$ reactive monomers carrying a B molecule with probability

$$K(N_1 \to N_1 - 1) = \frac{1}{N_1}\,. \tag{10}$$

The detailed balance condition follows

$$\pi_{N_1} K(N_1 \to N_1 - 1)\text{acc}(N_1 \to N_1 - 1) = \pi_{N_1+1} K(N_1 + 1 \to N_1)\text{acc}(N_1 + 1 \to N_1) \tag{11}$$

where acc() is the acceptance probability and $\pi$ the Boltzmann distribution (see Main Eqs. 1, 2). Using the first equality of Main Eq. 6 in the Metropolis acceptance rule, we obtain

$$\text{acc}(N_1 \to N_1 + 1) \;=\; \min\left[1, \frac{N_\text{R} - N_1 - 2N_2}{N_1 + 1} K_a \rho_B \exp[-\beta\, \Delta U_\text{conf}]\right], \tag{12}$$

$$\text{acc}(N_1 \to N_1 - 1) \;=\; \min\left[1, \frac{N_1}{N_\text{R} - N_1 - 2N_2 + 1} \frac{1}{K_a \rho_B} \exp[-\beta\, \Delta U_\text{conf}]\right] \tag{13}$$

$\Delta U_\text{conf}$ is the change in configurational energy calculated using Main Eq. 7. Given that this MC move does not update the chain backbone ($\{r'\}$=$\{r\}$), $\Delta U_\text{conf} \neq 0$ only in systems with IBMs given that, in this case, R monomers carrying or not a B molecule interact differently.

### 2.3 Loop/Unloop MC moves

In this section, we describe the MC move that, with equal probability, attempts either to form or open a loop.

**Forming a loop.** We first report the procedure to generate a looped configuration $\Gamma^{R_2B}_\text{new}$ (see right panel of Fig. S7a), starting from a complementary pair of R monomers ($i$ and $j$ in the left panel of Fig. S7a). By complementary pair, we mean that only one of the two R monomers ($i$ or $j$) carries a B molecule. We start by selecting a reactive monomer $i$ not

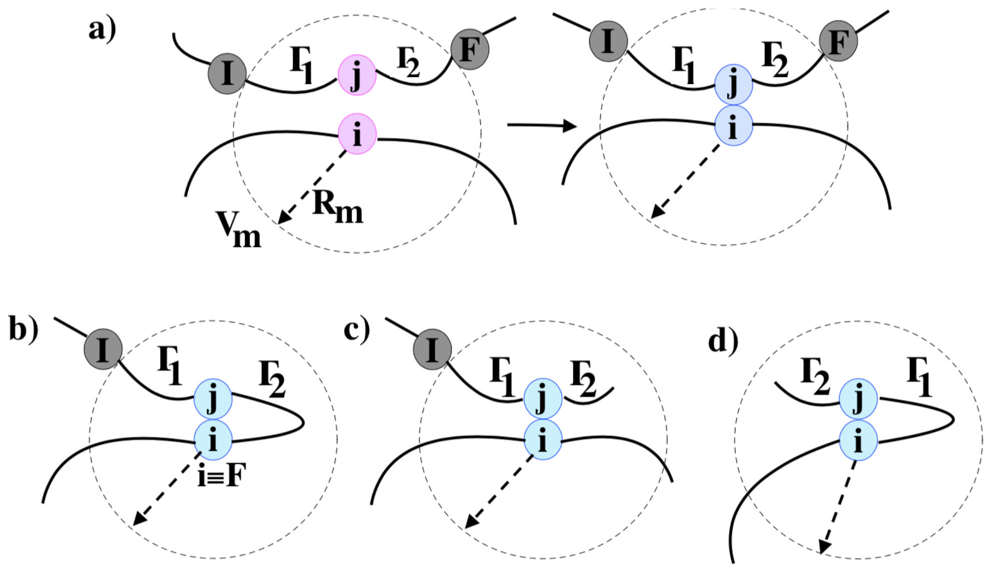


Figure S7: **a)** Loop formation between two complementary monomers $i$ and $j$. Monomer $i$ defines the center of the sphere $V_m$ of radius $R_m$, and $j$ is found inside this sphere. The chain section $\Gamma$ (limited by monomers $I$ and $F$) is reconfigured to form a loop by reacting $i$ with $j$. **b-d**) Different types of chain segments. **b**) The final monomer $F$ coincides with monomer $i$. **c**) The terminal of $\Gamma$ coincides with the end monomer of the polymer. $F$ does not exist and $\Gamma_2$ is (re)grown without constraining the position of the endpoint (see Eq. 8) **d**) The $F$ monomer coincides with $i$ (as in panel **b**) and $\Gamma_2$ includes the end of the chain (as in panel **c**). In all cases, $I$ and $F$ are the same in the bound and unbound state. $\Gamma$ comprises two segments ($\Gamma_1$ and $\Gamma_2$) and monomer $j$.

forming a loop with probability

$$P_i = \frac{1}{N_R - 2 \cdot N_2} \, . \tag{14}$$

We then select a monomer $j$ within the $\bar{n}_1$ R monomers complementary to $j$ found in a sphere ($V_m$) of radius $R_m$ centered on $i$ with probability

$$P_j = \frac{1}{\bar{n}_1} \, . \tag{15}$$

The sphere $V_m$ defines the proximity space where a reaction between $i$ and $j$ can take place. The radius of the sphere $R_m$ does not affect the results; it solely determines the efficiency of the algorithm [5]. In the present work, we have fixed $R_m$ to $R_m = 10\sigma$. The combined probability of selecting a pair of complementary monomers ($i$ and $j$) at a given connectivity state $(\nu_1, \nu_2)$ is given by

$$P_{\text{comb}}(i, j|\nu_1, \nu_2) = \frac{1}{(N_R - 2 \cdot N_2)\bar{n}_1} \tag{16}$$

Given $j$, we determine the longest segment $\Gamma$ ($\Gamma = \Gamma_1 \cup \mathbf{r}_j \cup \Gamma_2$ in Fig. S7) included in $V_m$ and containing $j$. In general, $\Gamma$ is limited by monomers $I$ and $F$ located just outside $V_m$ (Fig. S7a and Fig. S7b). However, if $\Gamma_1$ ($\Gamma_2$) carries monomers forming loops, $I$ ($F$) coincides with the looped monomer that is closest to $j$. $I$ or $F$ may also coincide with $i$ (as in Fig. S7b and Fig. S7d). The monomers $I$ and $F$ are determined by moving along the chain from $i$ towards $j$. $F$ does not exist when $\Gamma$ contains one of the polymer ends (as in Fig. S7c and Fig. S7d). In the latter case, the growth of $\Gamma_2$ is not biased by the guiding function $p(N, r)$. Instead, the trials bonds $\mathbf{u}_i^n$ are sampled from a uniform distribution as discussed in Sec. 2.1.

When generating a configuration with reacted $i$ and $j$ (in the following labelled with $\Gamma_{\text{new}}^{R_2B}$), we first delete the existing segment of chain $\Gamma$ from the system and then insert monomer $j$ around $i$ at a distance equal to one bond length $\sigma$ by selecting $\mathbf{r}_j$ from a list of trials. Importantly, if $j = I + 1$ or $j = F - 1$, the trials are generated using a crankshaft move (see Sec. 2.1). Instead, if $j > I + 1$ and $j < F - 1$, the trials are generated over the sphere $\Omega_0$ centered on $i$. The distribution of the trials ($p_{\text{trials}}$) comprises guiding functions $p$ pulling the position of monomer $j$ towards $F$ and $I$ [6]

$$p_{\text{trials}} = \begin{cases} \frac{p(N_b+1, r_{j,F})}{2\pi Z_j^i} & Z_j^i = \frac{\sum_{n=1}^{N_\alpha} p(N_b+1, r_{j,F}^n)}{N_\alpha} \quad \text{if } j = I + 1 \\ \frac{p(N_a+1, r_{I,j})}{2\pi Z_j^i} & Z_j^i = \frac{\sum_{n=1}^{N_\alpha} p(N_a+1, r_{I,j}^n)}{N_\alpha} \quad \text{if } j = F - 1 \\ \frac{p(N_a+1, r_{I,j}) p(N_b+1, r_{j,F})}{\Omega_0 Z_j^i} & Z_j^i = \frac{\sum_{n=1}^{N_\alpha} p(N_a+1, r_{I,j}^n) p(N_b+1, r_{j,F}^n)}{N_\alpha} \quad \text{if } I + 1 < j < F - 1 \end{cases} \tag{17}$$

where $N_a$ and $N_b$ are the number of monomers entering the fraction of $\Gamma$ between $j$ and, respectively, $F$ and $I$. $Z_j^i$ are unbiased estimations of the normalization of $p_{\text{trials}}$ that are calculated using $N_\alpha$ random positions ($\mathbf{r}_j^n$) uniformly distributed on the sphere centered on

$i$. The selection is implemented using a probability distribution ($p_{\text{select}}$), which is similar to what used in Sec. 2.1

$$p_{\text{select}} = \begin{cases} \frac{\exp[-\beta\, u_j^n]}{W_j} & \text{if } j = I+1 \\ \frac{\exp[-\beta\, u_j^n]}{W_j} & \text{if } j = F-1 \\ \frac{\exp[-\beta\, u_B^n]}{W_j} & I+1 < j < F-1 \end{cases} \tag{18}$$

where we introduced the Rosenbluth factors $W_j$ defined as in Sec. 2.1. Following the procedure described above, the position of monomer $j$ is generated with probability

$$P_{\text{grow}}(\mathbf{r}_j) = \delta(|\mathbf{r}_i - \mathbf{r}_j| - \sigma) \cdot p_{\text{trials}} \cdot p_{\text{select}}\,. \tag{19}$$

After inserting monomer $j$, we grow $\Gamma_1$ and $\Gamma_2$ (in the following $\Gamma_{\text{new}}^{R_2B(I,j)}$ and $\Gamma_{\text{new}}^{R_2B(j,F)}$) sequentially using the methods of Sec. 2.1. Importantly, the statistical weights of trials found outside the reacting sphere $V_m$ is taken equal to zero [5]. The probability of growing $\Gamma_{\text{new}}^{R_2B}$ and the associated Rosenbluth factors are then given by (when growing $\Gamma_{\text{new}}^{R_2B(I,j)}$ and then $\Gamma_{\text{new}}^{R_2B(j,F)}$)

$$\begin{aligned} P_{\text{grow}}(\Gamma_{\text{new}}^{R_2B}) &= P_{\text{grow}}(\mathbf{r}_j) \cdot P_{\text{grow}}(\Gamma_{\text{new}}^{R_2B(I,j)}|\mathbf{r}_j) \cdot P_{\text{grow}}(\Gamma_{\text{new}}^{R_2B(j,F)}|\Gamma_{\text{new}}^{R_2B(I,j)}, \mathbf{r}_j) & (20) \\ W_{\Gamma_{\text{new}}^{R_2B}} &= W_j \cdot W_{\Gamma_{\text{new}}^{R_2B(I,j)}} \cdot W_{\Gamma_{\text{new}}^{R_2B(j,F)}} & (21) \end{aligned}$$

If $j = I+1$ ($j = F-1$), then $P_{\text{grow}}(\Gamma_{\text{new}}^{R_2B(I,j)})$ ($P_{\text{grow}}(\Gamma_{\text{new}}^{R_2B(j,F)})$) is taken equal to one along with the corresponding Rosenbluth weight. The Rosenbluth factors depend on the order in which $\Gamma_{\text{new}}^{R_2B(I,j)}$ and $\Gamma_{\text{new}}^{R_2B(j,F)}$ are grown as the interactions between the two segments enter into the Rosenbluth factor relative to the second insertion. Finally, the probability ($P_{\text{gen}}$) of generating a new segment forming a loop $\Gamma^{R_2B}_{\text{new}}$ reads as

$$P_{\text{gen}}(\Gamma^{R_2B}_{\text{new}}) = P_{\text{comb}}(i,j|\nu_1,\nu_2)P_{\text{grow}}(\Gamma^{R_2B}_{\text{new}}). \tag{22}$$

The probability $P_{\text{gen}}$ of generating an old configuration $\Gamma^{R_2B}_{\text{old}}$ is estimated as done in Configurational Biased Monte Carlo (CBMC) using $k-1$ trial segments to regrow $\Gamma^{R^2B}_{\text{old}}$ (starting from $\mathbf{r}_j$) while selecting the existing monomer (see Sec. 2.1). The expression of $P_{\text{gen}}(\Gamma^{R_2B}_{\text{old}})$ follows from Eq. 22.

**Opening a loop.** We now describe how we unloop an already reacted pair of monomers $i$ and $j$. We first randomly select a pair of cross-linked monomers with a probability given by

$$P_{\text{comb}}(ij|\nu_1,\nu_2) = \frac{1}{2N_2} \tag{23}$$

where the factor 1/2 is the probability of centering the reacting sphere $V_m$ on one of the two R monomers. After identifying $i$ and $j$, we calculate $I$ and $F$ as done above (see Fig. S7) and generate a new free configuration $\Gamma_{\text{new}}$ constrained to lay inside $V_m$ between $I$ and $F$. Then

$$P_{\text{gen}}(\Gamma_{\text{new}}) \quad = \quad P_{\text{comb}}(ij|\nu_1,\nu_2)\cdot P_{\text{grow}}(\Gamma_{\text{new}}) \tag{24}$$

where $P_{\text{comb}}$ is given by Eq. 23 and $P_{\text{grow}}$ by Eq. 6 in which the Rosenbluth weights do not include trial positions outside $V_m$. Accordingly, we calculate the probability of regrowing an existing configuration not forming a loop ($\Gamma_{\text{old}}$) using Eq. 24.

**Acceptance probabilities.** $P_{\text{gen}}(\Gamma)$, $P_{\text{gen}}(\Gamma^{R_2B})$, and the Boltzmann weights of the new/old configurations ($\pi_{\text{new}} = \pi(\Gamma^{R_2B}_{\text{new}},\nu'_1,\nu'_2)$ and $\pi_{\text{old}} = \pi(\Gamma_{\text{old}},\nu_2)$, where $\nu_1/\nu'_1$ and $\nu_2/\nu'_2$ are the connectivity states of the old/new configuration) are then used to determine the acceptance rate of the proposed and reverse move, $\text{acc}(\Gamma_{\text{old}} \to \Gamma^{R_2B}_{\text{new}})$ and $\text{acc}(\Gamma^{R_2B}_{\text{old}} \to \Gamma_{\text{new}})$. Using the detailed balance conditions [4]

$$\pi_{\text{old}} P_{\text{gen}}(\Gamma^{B_2R}_{\text{new}}) \text{acc}(\Gamma_{\text{old}} \to \Gamma^{B_2R}_{\text{new}}) = \pi_{\text{new}} P_{\text{gen}}(\Gamma_{\text{old}}) \text{acc}(\Gamma^{B_2R}_{\text{new}} \to \Gamma_{\text{old}}) \tag{25}$$

$$\pi_{\text{old}} P_{\text{gen}}(\Gamma_{\text{new}}) \text{acc}(\Gamma^{B_2R}_{\text{old}} \to \Gamma_{\text{new}}) = \pi_{\text{new}} P_{\text{gen}}(\Gamma^{B_2R}_{\text{old}}) \text{acc}(\Gamma_{\text{new}} \to \Gamma^{B_2R}_{\text{old}}) \tag{26}$$

along with Eqs. 6, 8, and Main Eqs. 5, we can write the Metropolis acceptance rates of binding and unbinding two complementary monomers as follows

$$\text{acc}(\Gamma_{\text{old}} \to \Gamma^{R_2B}_{\text{new}}) = \min\left[1, K_a \frac{(N_R - 2 \cdot N_2)\overline{n}_1}{N_2 + 1} \frac{P_{\text{grow}}(\Gamma_{\text{old}}|\overline{\Gamma})}{\Omega_0 P_{\text{grow}}(\Gamma^{R_2B}_{\text{new}}|\overline{\Gamma})} \exp(-\beta \Delta U)\right] \tag{27}$$

$$\text{acc}(\Gamma^{R_2B}_{\text{old}} \to \Gamma_{\text{new}}) = \min\left[1, \frac{1}{K_a} \frac{N_2}{((N_R - 2 \cdot N_2) + 1)(\overline{n}_1 + 1)} \frac{P_{\text{grow}}(\Gamma^{R_2B}_{\text{old}}|\overline{\Gamma})}{\Omega_0 P_{\text{grow}}(\Gamma_{\text{new}}|\overline{\Gamma})} \exp(-\beta \Delta U)\right] \tag{28}$$

where $\Delta U$ is the change in configurational energy given by $U_{\nu_1,\nu_2}$ (Main Eq. 2). By using Eq. 6, we get

$$\frac{P_{\text{grow}}(\Gamma_{\text{old}})}{P_{\text{grow}}(\Gamma^{R_2B}_{\text{new}})} = \frac{W_{\Gamma^{R_2B}_{\text{new}}} \cdot Z^i_j \cdot \Omega_0 \cdot \exp(\beta \Delta U)}{W_{\Gamma_{\text{old}}} \cdot p(N_a + N_b + 2, r_{I,F})} \tag{29}$$

with $Z^i_j$ given by Eq. 17. The final expressions of the acceptance rates are then

$$\text{acc}(\Gamma_{\text{old}} \to \Gamma^{R_2B}_{\text{new}}) = \min\left[1, K_a \frac{(N_R - 2 \cdot N_2) \cdot \bar{n}_1}{(N_2 + 1)} \frac{Z^i_j}{p(r_{\bar{\Gamma}}, N_a + N_b + 2)} \frac{W_{\Gamma^{R_2B}_{new}}}{W_{\Gamma_{old}}}\right] \tag{30}$$

$$\text{acc}(\Gamma^{R_2B}_{\text{old}} \to \Gamma_{\text{new}}) = \min\left[1, \frac{1}{K_a} \frac{N_2}{(\bar{n}_1 + 1)((N_R - 2 \cdot N_2) + 1)} \frac{p(r_{\bar{\Gamma}}, N_a + N_b + 2)}{Z^i_j} \frac{W_{\Gamma_{new}}}{W_{\Gamma^{R_2B}_{old}}}\right] \tag{31}$$

### 2.4 Loop/Unloop+Unbind/Bind MC moves

In this section, we describe the MC move that, with equal probability, attempts either to form a loop while detaching a B molecule or to open a loop while binding a B molecule to the chain (see Main Fig. 3b). The (re)growth of open/bound segments follows the same procedure of the previous section, while the selection of the reacting monomers ($i$ and $j$) changes. In particular, when attempting to form a loop, we first select $i$ within the $N_1$ monomers carrying a B molecule and then select $j$ within the $\overline{n}_1$ monomers carrying a B molecule in the sphere $V_m$ centered on $i$. Using the notation of the previous section, the acceptance probabilities of the loop and unloop move are given by

$$\mathrm{acc}(\Gamma_{\mathrm{old}}^{RB+RB} \to \Gamma_{\mathrm{new}}^{R_2B}) = \min\left[1, \frac{1}{\rho_B} \frac{N_1 \ \cdot \bar{n}_1}{(N_2+1)} \frac{Z_j^i}{p(r_{\bar{\Gamma}}, N_i + N_j + 2)} \frac{W_{\Gamma_{\mathrm{new}}^{R_2B}}}{W_{\Gamma_{\mathrm{old}}}}\right] \tag{32}$$

$$\mathrm{acc}(\Gamma_{\mathrm{old}}^{R_2B} \to \Gamma_{\mathrm{new}}^{RB+RB}) = \min\left[1, \rho_B \frac{N_2}{(N_1+2)(\bar{n}_1+1)} \frac{p(r_{\bar{\Gamma}}, N_i + N_j + 2)}{Z_j^i} \frac{W_{\Gamma_{\mathrm{new}}}}{W_{\Gamma_{\mathrm{old}}^{R_2B}}}\right] \tag{33}$$

Notice that in the denominator of the r.h.s. of Eq. 33 we have $N_1 + 2$ as, after opening a loop, the number of monomers carrying a B molecules and not forming a loop increases by two (see Main Fig. 3b). $1/\rho_B$ and $\rho_B$ are the entropic contributions, respectively, of detaching or attaching a B molecule to the chain as in Eqs. 12, 13. However, as compared to the acceptance probabilities of the bind/unbind MC move, Eqs. 35, 33 are not a function of $K_a$ as the number of reacted $R$ monomers is not changed by the move ($N_1 + 2N_2$ stays constant).

### 2.5 Swap MC move

In the swap MC move, a pair of reacted monomers ($ik$) is selected along with a third R monomer $j$ from all available free reactive sites within the sphere $V_m$ centered on $i$. As in the loop/unloop move (Sec. 2.3), $k$ and $j$ define two segments (that may coincide [6]) limited by two pairs of monomers ($k \in [I_k,\, F_k]$ and $j \in [I_j,\, F_j]$). We detach monomer $k$ from $i$ and grow a configuration in which monomer $j$ forms a loop with $i$ (See Main Fig. 3b). The new configuration is accepted with probability

$$\mathrm{acc}(\Gamma_{\mathrm{old}}^{I_k,i,F_k} \to \Gamma_{\mathrm{new}}^{I_j,i,F_j}) \;=\; \min\left[1, \frac{Z_k^i}{Z_j^i}\, \frac{W_{\Gamma_{\mathrm{new}}^{I_j,i,F_j}}}{W_{\Gamma_{\mathrm{old}}^{I_k,i,F_k}}} \frac{W_{\Gamma_{\mathrm{new}}^{I_k,F_k}}}{W_{\Gamma_{\mathrm{old}}^{I_j,F_j}}}\right], \tag{34}$$

where $\Gamma^{I_k,i,F_k}$ ($\Gamma^{I_j,i,F_j}$) is the $[I_k,\, F_k]$ ($[I_j,\, F_j]$) segment in which $k$ ($j$) is bound to $i$, while $\Gamma^{I_k,F_k}$ ($\Gamma^{I_j,F_j}$) is the $[I_k,\, F_k]$ ($[I_j,\, F_j]$) segment without loops. The Rosenbluth weights, $Z_k^i$, and $Z_j^i$ are calculated as in Sec. 2.3. If both monomers $j$ and $k$ belong to the same chain section ($I_k = I_j$ and $F_k = F_j$), the swap MC move corresponds to a slide move of the segment containing $j$ and $k$ along the segment supporting $i$. In this case, we never (re)grow free segments ($\Gamma^{I_k,F_k}$ or $\Gamma^{I_j,F_j}$) and the corresponding Rosenbluth weights in Eq. 34 are taken equal to 1 ($W_{\Gamma_{\mathrm{new}}^{I_k,F_k}} = W_{\Gamma_{\mathrm{old}}^{I_j,F_j}} = 1$).

### 2.6 Swing MC move

In the swing MC move, we randomly choose two pairs of reacted monomers, $(ij)$ and $(kl)$ ($(kl)$ inside the reacting sphere $V_m$ centered on $i$), and propose a new configuration in which $l$ and $j$ binds, respectively, $i$ and $k$ (see Main Fig. 3b). This is done by simultaneously reconfiguring the segments supporting $l$ and $j$ ($\Gamma_{\mathrm{old}}^{I_l,k,F_l}$ and $\Gamma_{\mathrm{old}}^{I_j,i,F_j}$, see Sec. 2.5). Following the considerations of the previous sections we write the acceptance probability as

$$\mathrm{acc}(\{\Gamma_{\mathrm{old}}^{I_l,k,F_l}, \Gamma_{\mathrm{old}}^{I_j,i,F_j}\} \to \{\Gamma_{\mathrm{new}}^{I_l,i,F_l}, \Gamma_{\mathrm{new}}^{I_j,k,F_j}\}) \;=\; \min\left[1, \frac{W_{\Gamma_{\mathrm{new}}^{I_l,i,F_l}} W_{\Gamma_{\mathrm{new}}^{I_j,k,F_j}} Z_j^i Z_l^k}{W_{\Gamma_{\mathrm{old}}^{I_l,k,F_l}} W_{\Gamma_{\mathrm{old}}^{I_j,i,F_j}} Z_i^l Z_k^j}\right]. \tag{35}$$

[11] H Yamakawa. *Modern theory of polymer solutions*. Harper & Row, 1971.
